# Stabilization of Integrator/INTAC by the small but versatile DSS1 protein

**DOI:** 10.1101/2024.02.25.581915

**Authors:** Congling Xu, Qian-Xing Zhou, Hai Zheng, Aixia Song, Wen-Ying Zhao, Yan Xiong, Zixuan Huang, Yanhui Xu, Jingdong Cheng, Fei Xavier Chen

## Abstract

Integrator-PP2A (INTAC) is a highly modular complex orchestrating the transition of paused RNA polymerase II into productive elongation or promoter-proximal premature termination, with its loss resulting in transcription dysregulation and genome instability. Here, we identify human DSS1—a flexible 70-residue protein found in multiple functionally diverse complexes including the 26S proteasome—as an integral subunit of the INTAC backbone. Structural analysis of DSS1–INTAC, both alone and in association with paused polymerase, demonstrates intimate interactions between DSS1 and the INTAC backbone subunit INTS7. We identify tryptophan 39 of DSS1 as being critical for interacting with INTAC and find that mutating this residue disrupts DSS1’s interaction with INTAC, while maintaining DSS1’s interaction with the proteasome. This substitution not only destabilizes INTAC and therefore INTAC-dependent transcriptional regulation, but also reveals that INTAC is DSS1’s major chromatin-bound form. Together, our findings reveal the essential role of the versatile DSS1 protein in the structure and regulatory functions of INTAC.

## Abstract

Transcription by RNA Polymerase II (Pol II) in metazoans is a complex and finely tuned process that includes a series of phases, starting with initiation, moving through promoter-proximal pausing and release, progressing to productive elongation, and culminating in termination (*1–4*). The process begins with the assembly of the pre-initiation complex (PIC) at promoters, where general transcription factors recruit Pol II, leading to the melting of double-stranded DNA and synthesis of a nascent transcript (*5–7*). Pol II then enters a promoter-proximal paused state within 200 bp downstream of the transcription start sites (TSSs). After pausing, Pol II can either progress to productive elongation for RNA synthesis or be terminated prematurely, making way for the subsequent transcription cycle (*8–11*).

Promoter-proximal pausing of Pol II is stabilized by the negative elongation factor (NELF) and DRB sensitivity-inducing factor (DSIF), with NELF anchoring Pol II near its pausing sites and DSIF preventing premature termination and proteolysis of Pol II subunits (*12–20*). The transition into elongation is predominantly facilitated by P-TEFb composed of kinase CDK9 and a regulatory subunit cyclin T1, which phosphorylates Pol II, NELF, and DSIF among its targets (*1, 2, 21, 22*). Conversely, the decision for premature termination of promoter-proximal Pol II is significantly influenced by the recently identified Integrator-PP2A (INTAC) complex (*23–25*).

The multisubunit INTAC complex, with its dual catalytic activities, is composed of six distinct modules. Among these modules, the backbone and shoulder modules provide a structural framework, enabling the positioning of the RNA endonuclease and phosphatase modules on opposite sides of this scaffold (*25*). This structural arrangement allows the decoupling of the two catalytic modules, which exert differential impacts on the fate of paused Pol II and gene expression (*26*). The SOSS module, featuring the single-stranded DNA (ssDNA)-binding protein SSB1/2, targets ssDNA within R-loops, thereby facilitating the recruitment of INTAC to R-loop-enriched genomic regions (*27*). The auxiliary/arm module of INTAC mediates the recognition of transcription factors, and it has some INTAC-independent roles in the regulation of transcription (*28–31*).

Integrator was initially identified through the purification of deleted in split hand/split foot 1 protein (DSS1) by Shiekhattar and colleagues two decades ago (*32*). DSS1, a homolog of suppressor of exocyst mutations 1 (SEM1) from yeast, is characterized by its small size and flexibility, being intrinsically disordered on its own (*33, 34*). In addition to being a potential component of Integrator and INTAC, DSS1/SEM1 also associates with several other complex assemblages, including the 26S proteasome and TREX-2 in yeast and mammals, and additionally in mammals with BRCA2 complexes (*34–39*). Within the proteolysis pathway, DSS1/SEM1 is not only integrated into the 19S regulatory complex, contributing to the efficient assembly of the 26S proteasome, but it might also act as a ubiquitin receptor (*35, 36, 40, 41*). The incorporation of DSS1/SEM1 into TREX-2 or BRCA2 occurs independently of its association with the 26S proteasome. In the TREX-2 complex, DSS1 directly binds and stabilizes PCID2, which facilitates the formation of a platform for mRNA binding and export from the nucleus (*37, 38, 42*). Moreover, as an integral and conserved partner of the BRCA2 complex, DSS1 is crucial for the conformational dynamics and stability of BRCA2, ensuring its function in homologous recombination repair of double-strand DNA breaks (DSBs) (*39, 43, 44*). Despite the well-defined functions of DSS1 in some of its complexes, the nature of its interaction with Integrator/INTAC, as well as the extent to which it functions in the steps of transcriptional regulation executed by this complex, have been largely unexplored.

In this study, we highlight DSS1 as an integral component of the INTAC backbone module, essential for maintaining the stability of INTAC and thus enabling its diverse functions in transcription regulation.

## Results

### DSS1 associates with the INTAC complex

To determine whether DSS1 genuinely constitutes a component of the Integrator or INTAC complex, we engineered a cell line with inducible expression of Flag-tagged DSS1 using the HEK293 Flp-In T-REx system, followed by Flag-affinity purification and mass spectrometry (Fig. 1A). Our analysis of the proteomics data revealed the enrichment of all INTAC subunits, including phosphatase module subunits PP2A-A and PP2A-C, the presence of which distinguishes INTAC from Integrator alone (Fig. 1B) (*45*). Further analysis also retrieved several known DSS1-containing complexes, such as the 26S proteasome, TREX-2, and BRCA2 complexes (Fig. 1C). To verify the endogenous interaction between DSS1 and INTAC, we integrated a Flag tag, along with a FKBP12^F36V^ degradation tag (dTAG) for rapid depletion, at the endogenous *DSS1* locus in the human colorectal cancer cell line DLD-1 (Fig. 1D). Co-immunoprecipitation with a Flag antibody in this modified DSS1–Flag–dTAG cell line successfully co-purified all examined INTAC subunits across different modules (Fig. 1E). Reciprocally, endogenous immunoprecipitation experiments targeting INTS3 and INTS5 further substantiated their association with DSS1 (Fig. 1F). These data support a close physical association of DSS1 with INTAC.

**Fig. 1.**
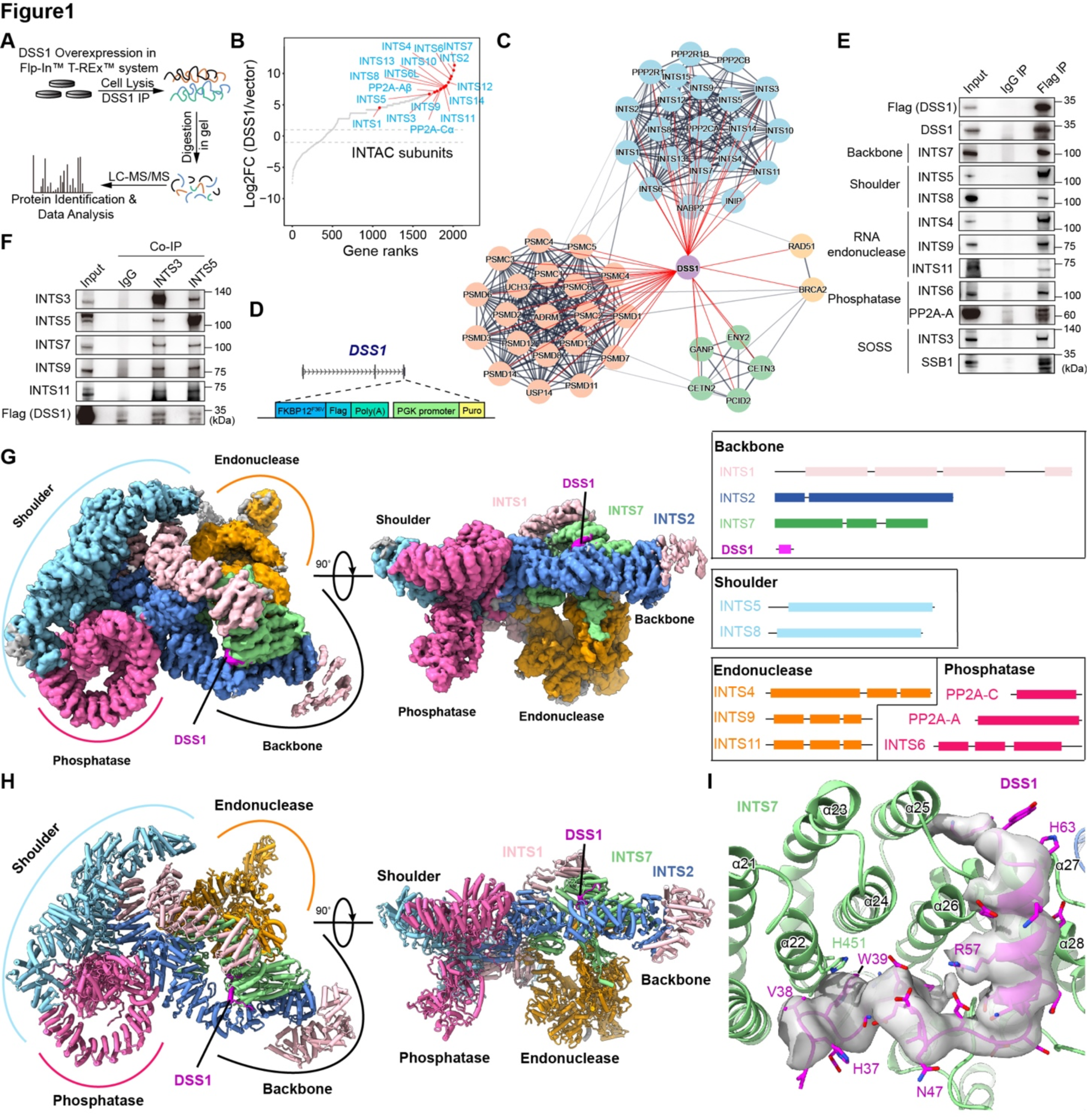
DSS1 is an integral component of the INTAC backbone module. (A) The workflow of DSS1 purification and mass spectrometry (MS) using Flp-In T-Rex system in HEK293 cells. (B) Proteins identified by MS were ranked by the enrichment in purification of DSS1 versus vector. INTAC subunits are labeled. (C) STRING protein-protein interaction network of DSS1 interactome combined with the proteomics data of DSS1. Red lines indicate the interactions identified in our DSS1 MS data. (D) Schematic of the generation of DSS1–Flag–dTAG cells. (E) Co-IP analysis of Flag (DSS1) followed by western blotting in DSS1–Flag–dTAG cells. (F) Co-IP analysis of endogenous INTS3 and INTS5 in DSS1–Flag–dTAG cells. (G-H) Overall cryo-EM map (G) and the molecular model (H) of the DSS1-INTAC complex in two different views, with subunit surfaces/models colored as schematic module (right of panel G). Notably, the cryo-EM map is a composite map derived from multibody refinement. (I) A close-up view of DSS1 (magenta) in DSS1-INTAC was shown. DSS1 is represented as sticks, surrounded by a transparent density map, indicating its nice fitting.

### Structure of a DSS1-containing INTAC complex

To elucidate the molecular mechanisms of the interaction between DSS1 and INTAC, we determined the structure of the reconstituted DSS1–INTAC complex using cryo-electron microscopy (cryo-EM) single-particle analysis. The cryo-EM map was refined to an overall resolution of 4.1 Å, with the maps of each module improved to a resolution ranging from 3.7 to 4.2 Å by multibody refinement (Fig. 1G, and fig. S1, and table S1). A structural model was built by fitting previously determined structures of INTAC (*45*) into the cryo-EM maps followed by manual adjustment (Fig. 1H, and fig. S1, and table S1).

In our DSS1–INTAC structure, the backbone and shoulder modules of INTAC establish a cruciform scaffold, positioning the RNA endonuclease and phosphatase modules on opposite sides (Fig. 1, G and H), which is consistent with previously described INTAC configurations (*45–47*). The SOSS and auxiliary modules of INTAC, noted for their dynamic nature (*27, 48*), were not captured in our visualization. Notably, we identified an additional density within the backbone module closely interacting with INTS7. This allowed us to model the central and C-terminal segments of DSS1, spanning alanine 36 to methionine 67, into our structure (Fig. 1, G and H). The association of DSS1 and INTS7 is characterized by extensive hydrophobic interactions, situating DSS1 within a groove formed by α-helices of INTS7 (Fig. 1I). Furthermore, DSS1’s placement is secured by the arch-shaped structure of INTS2, which likely restricts DSS1’s exposure to external proteins (Fig. 1, G to I). These structural insights suggest that DSS1 is an integral component of the INTAC backbone module.

In addition to the DSS1–INTAC structure presented in this study, structures of DSS1 within the 26S proteasome, TREX-2, and BRCA2 complexes have been previously characterized (*34–39*). To understand the molecular mechanism underlying the multifaceted functions of DSS1, we compared its configurations across these different complexes. DSS1 presents strikingly unique structural adaptations in each context (fig. S2, A to D). While DSS1 tends to adopt α-helical structures at different positions of its C terminus in the various contexts, its N-terminal and central regions display remarkable flexibility, allowing for diverse structural accommodations to fit the interfaces of its interacting partners (fig. S2, A to D). This structural compliance of DSS1 could underly its ability to perform analogous scaffolding roles in otherwise unrelated complexes that are implicated in various biological processes.

### The genomic distribution of DSS1 is similar to that of INTAC

We next sought to delineate the cellular distribution of DSS1. Time-course treatment with FKBP12^F36V^ degrader, dTAG^V^-1 (*49*), resulted in a marked depletion of endogenous DSS1 in DSS1–Flag–dTAG cells (Fig. 2A). Immunofluorescence analysis revealed that endogenous DSS1 can be localized to the nucleus, and the immunofluorescence signal was effectively eliminated upon dTAG treatment, confirming a nuclear pool of DSS1 (Fig. 2, B and C). We therefore asked whether DSS1 associates with chromatin-bound INTAC. Initial attempts of Flag ChIP-seq in DSS1–Flag–dTAG cells did not yield significant enrichment (data not shown), potentially due to crosslinking constraints that obscure antibody recognition by sequestering DSS1 within the hydrophobic pocket of the backbone module. Consequently, we utilized Flag CUT&Tag in DSS1–Flag–dTAG cells treated with DMSO or dTAG for 24 hours. This approach identified 54,757 DSS1-bound genomic regions in DMSO-treated cells, which were significantly decreased following dTAG treatment (Fig. 2, D and E). The control IgG samples showed markedly lower signal intensity, further validating the specificity of the Flag CUT&Tag signals (Fig. 2, D and E). These DSS1-associated genomic regions included promoters, as well as intergenic and intragenic regions (Fig. 2F), aligning with the distribution patterns of INTAC subunits (*24, 26, 50*). Among these regions, promoters exhibited the highest occupancy of DSS1 (Fig. 2G). Pearson correlation analysis demonstrated a strong correlation between the genomic localization of DSS1 and other INTAC subunits, as well as active chromatin markers of promoters and enhancers (fig. S2E). Moreover, DSS1 occupancy is comparable to that of other INTAC subunits, particularly at promoter regions that have the highest enrichment of INTAC (Fig. 2, G to I, and fig. S2F). Together, these results suggest a significant nuclear localization of DSS1 and a genomic distribution that is congruent with other INTAC subunits.

**Fig. 2.**
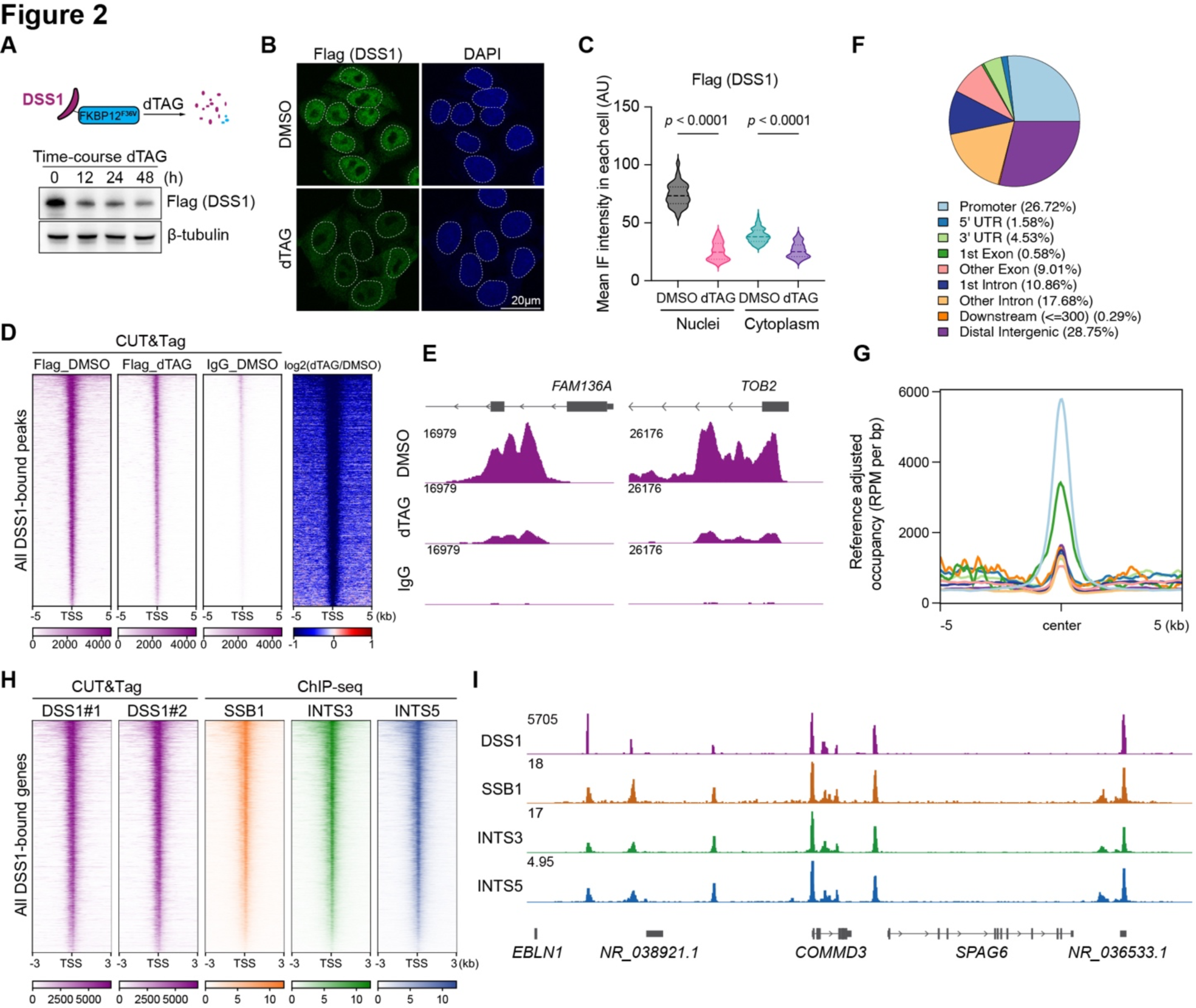
Similar genomic distribution of DSS1 and INTAC. (A) DSS1 degradation induced by time-course dTAG treatment in DSS1–Flag–dTAG cells. (B) Representative images showing the localization of endogenous DSS1 (green) along with the DAPI signal (blue) after 24-h DMSO or dTAG treatment in DSS1–Flag–dTAG cells. Data represent three independent experiments. (C) Quantification of mean immunofluorescence (IF) intensity of DSS1 in nuclei and cytoplasm of cells. (DMSO/nuclei: n = 110; dTAG/nuclei: n = 83; DMSO/cytoplasm, n = 111; dTAG/cytoplasm, n = 63). AU means arbitrary units. Outliners were excluded by ROUT method, Q = 10 %. P values were calculated using two-tailed unpaired t-tests. (D) Flag (DSS1) CUT&Tag signals over 10 kb regions centered on Flag (DSS1) peak summit in DSS1–Flag–dTAG cells treated with DMSO or dTAG for 24 h. (E) Representative track examples showing DSS1 occupancy in DSS1–Flag–dTAG cells treated with DMSO or dTAG for 24 h. (F) Pie chart showing genomic distributions of DSS1. (G) Metaplot of DSS1 occupancy on each DSS1-bound regions shown in (F). (H) Occupancy of DSS1, SSB1, INTS3, and INTS5 over 6 kb regions centered on the TSS of DSS1 target gene promoters. (I) Representative track examples showing the occupancy of DSS1, SSB1, INTS3, and INTS5.

### Selective disruption of the DSS1–INTAC interaction without affecting proteasomal DSS1

Our detailed structural analysis revealed significant hydrophobic interactions between DSS1 and the backbone subunit INTS7, which highlighted the importance of a tryptophan residue at position 39 (W39) within DSS1 for its association with INTAC (Fig. 3A and fig. S3, A and B). Immunoprecipitation assays demonstrated that substituting W39 in DSS1 with the positively charged arginine (W39R) significantly disrupted its interaction with INTAC without affecting its association with proteasome subunits such as PSMD4 and PSMD14 (Fig. 3B). Furthermore, mutations of residues within INTS7’s hydrophobic pockets similarly compromised INTAC’s interaction with DSS1 (fig. S3, C and D).

**Fig. 3.**
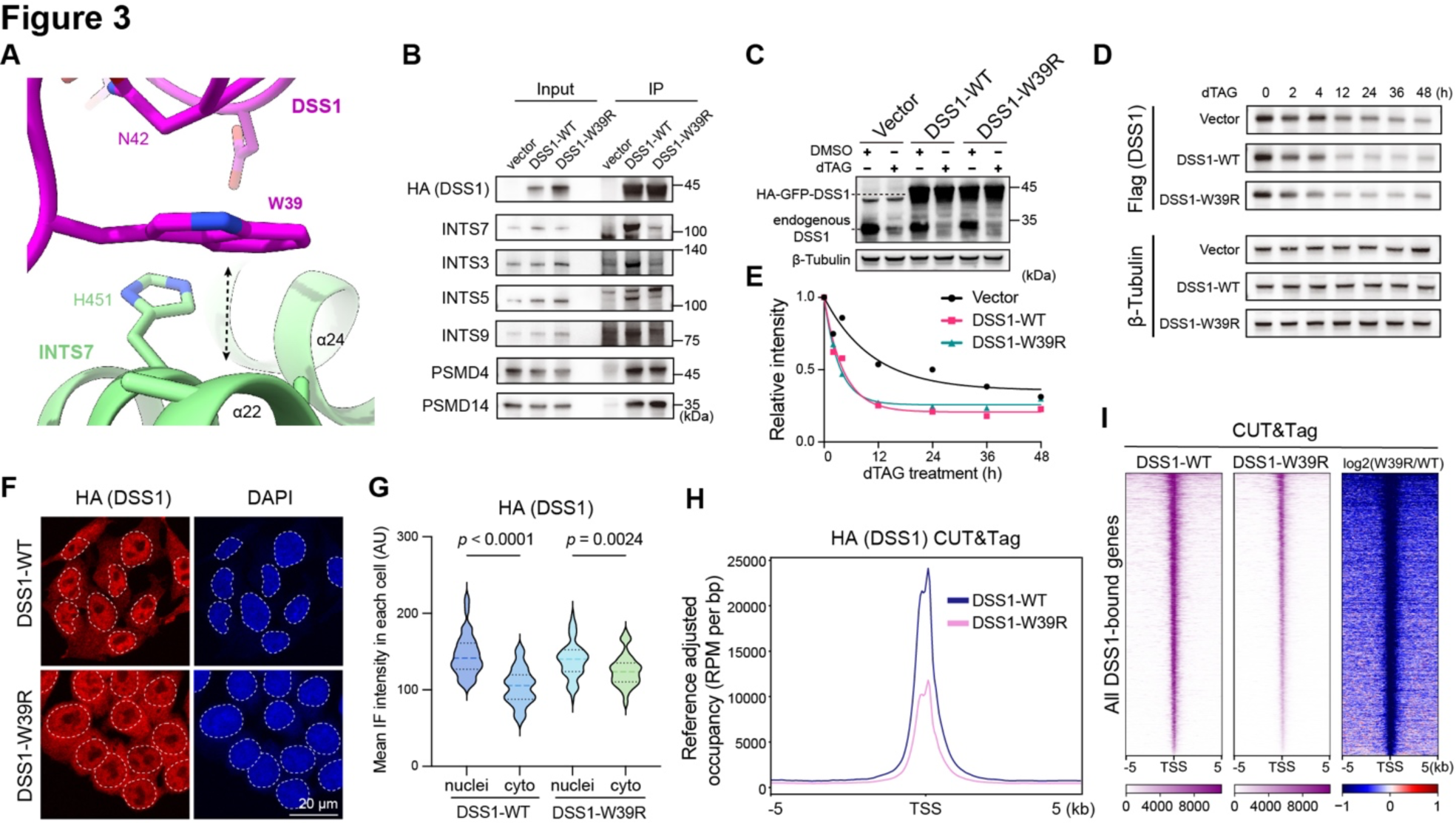
The DSS1–INTAC interaction is essential for the optimal chromatin recruitment of DSS1. (A) Detailed structural view of the interaction sites between INTAC and DSS1. Residue W39 stacks with the backbone of the helix 22 of INTS7, indicated by the bidirectional arrow. (B) Co-IP analysis of HA (DSS1) followed by western blotting. PSMD4 and PSMD14 are 26S proteasome subunits. (C) Overexpression of DSS1-WT, DSS1-W39R, or vector in DSS1–Flag–dTAG cells treated with DMSO or dTAG, followed by western blotting of DSS1. (D) Western blotting of Flag (DSS1) in DSS1-WT, DSS1-W39R, or vector overexpressing DSS1–Flag–dTAG cells with time-course dTAG treatment. (E) The quantified and normalized band intensities in (D). Exponential one phase decay model was utilized to fit nonlinear regression curve for Flag (DSS1) band intensities. (F) Representative images showing the localization of overexpressed DSS1-WT or DSS1-W39R (red) along with the DAPI signal (blue) after endogenous DSS1 degradation in DSS1–Flag–dTAG cells. Data represent two independent experiments. (G) Quantification of mean IF intensity of HA (DSS1) in nuclei and cytoplasm of DSS1–Flag–dTAG cells with DSS1-WT or DSS1-W39R. (WT/nuclei: n = 91; WT/cytoplasm: n = 90; W39R/nuclei, n = 82; W39R/cytoplasm, n = 82). AU means arbitrary units. Outliners were excluded by ROUT method, Q = 10 %. Multiple comparison after One-way-ANOVA was used to calculate p value. (H and I) Metaplot (H) and heatmaps (I) showing HA (DSS1) CUT&Tag signals over 10 kb regions centered on the TSS of target genes in dTAG-treated DSS1–Flag–dTAG cells expressing DSS1-WT or DSS1-W39R.

To assess the functional implications of the DSS1–INTAC interaction, we overexpressed HA–GFP-tagged wild-type (WT) DSS1 and the W39R mutant in DSS1–Flag–dTAG cells (Fig. 3C). Notably, the overexpression of either DSS1-WT or DSS1-W39R led to a more rapid degradation of the endogenous DSS1 compared to cells transfected with an empty vector (Fig. 3, D and E). This effect implies the involvement of DSS1 in supporting proteasomal activities independently of its association with INTAC. Supporting this notion, upon inhibition of protein synthesis with cycloheximide (CHX), the depletion of DSS1 extended the half-life of proteins typically subject to rapid turnover, such as CDC6 and C-MYC (fig. S3E) (*51*). Remarkably, reintroducing either DSS1-WT or the INTAC-interaction-deficient DSS1-W39R mutant restored the degradation rates of these proteins to their original levels (fig. S3E). These findings indicate that DSS1 is integrated into INTAC and the proteasome through distinct interfaces, with the W39 residue being crucial for its association with INTAC but dispensable for its role in proteasome-mediated degradation.

### The DSS1–INTAC interaction is required for the optimal chromatin recruitment of DSS1

Since perturbations of the proteasome could potentially impact the cellular levels of any protein and therefore have unpredictable and uninterpretable consequences, we compared the effects of expressing DSS1-WT and DSS1-W39R to differentiate the role of DSS1 within INTAC from its proteasomal activity. To this end, we expressed DSS1-WT or DSS1-W39R in DSS1–Flag–dTAG cells that were treated with dTAG to deplete endogenous DSS1. Immunofluorescence analysis revealed that DSS1-WT is predominantly localized to the nucleus, whereas DSS1-W39R was equally distributed across both nuclear and cytoplasmic compartments (Fig. 3, F and G). To evaluate their genomic distribution, we performed CUT&Tag for both DSS1-WT and DSS1-W39R in cells depleted for endogenous DSS1. Despite similar expression levels (Fig. 3C), DSS1-WT showed significantly higher occupancy on chromatin than DSS1-W39R on average (Fig. 3H). Heatmap analysis shows a widespread reduction in genomic occupancy for DSS1-W39R compared with DSS1-WT (Fig. 3I), which can be seen on example genes (fig. S3F). These findings indicate that the DSS1–INTAC interaction is required for the proper nuclear localization and chromatin association of DSS1.

### Structural analysis of the DSS1-containing INTAC–paused elongation complex (PEC)

The intimate interaction between DSS1 and INTAC seen in our structural studies, along with their similar genomic distributions, implies that DSS1 is likely present in most or all forms of INTAC. We and colleagues previously reported a structure of INTAC in association with the paused elongation complex (PEC) (*46, 47*), which typically is comprised of the paused form of Pol II, NELF and DSIF. Consistent with our hypothesis, purifications of DSS1 were enriched for subunits of Pol II, NELF and DSIF (fig. S4, A and B). Endogenous co-IP further corroborated the association of DSS1 with these PEC components (fig. S4C).

To better ascertain the incorporation of DSS1 into INTAC in the context of PEC, we reconstituted a complex containing DSS1–INTAC and PEC, followed by cryo-EM single-particle analysis. The resulting map reached an overall resolution of 4.6 Å, with individual sub-complexes improved to resolutions from 4.2 to 4.3 Å through multibody refinement. The structural model was fitted by integrating previously determined structures of the INTAC–PEC (*46*) into the cryo-EM maps, followed by manual adjustments (fig. S1 and table S1).

In our DSS1–INTAC–PEC structure, we observed an extra density corresponding to the N-terminus of INTS1, which is not visible in the standalone INTAC structure, in addition to the integration of the PEC components, including Pol II, NELF, and DSIF (Fig. 4, A and B). This “tail” of INTS1 extends from the bottom of the backbone module and loops back to interact with PEC. Overall, INTAC complex forms close contacts with PEC, featuring extensive interacting interfaces between them (Fig. 4, A and B).

**Fig. 4.**
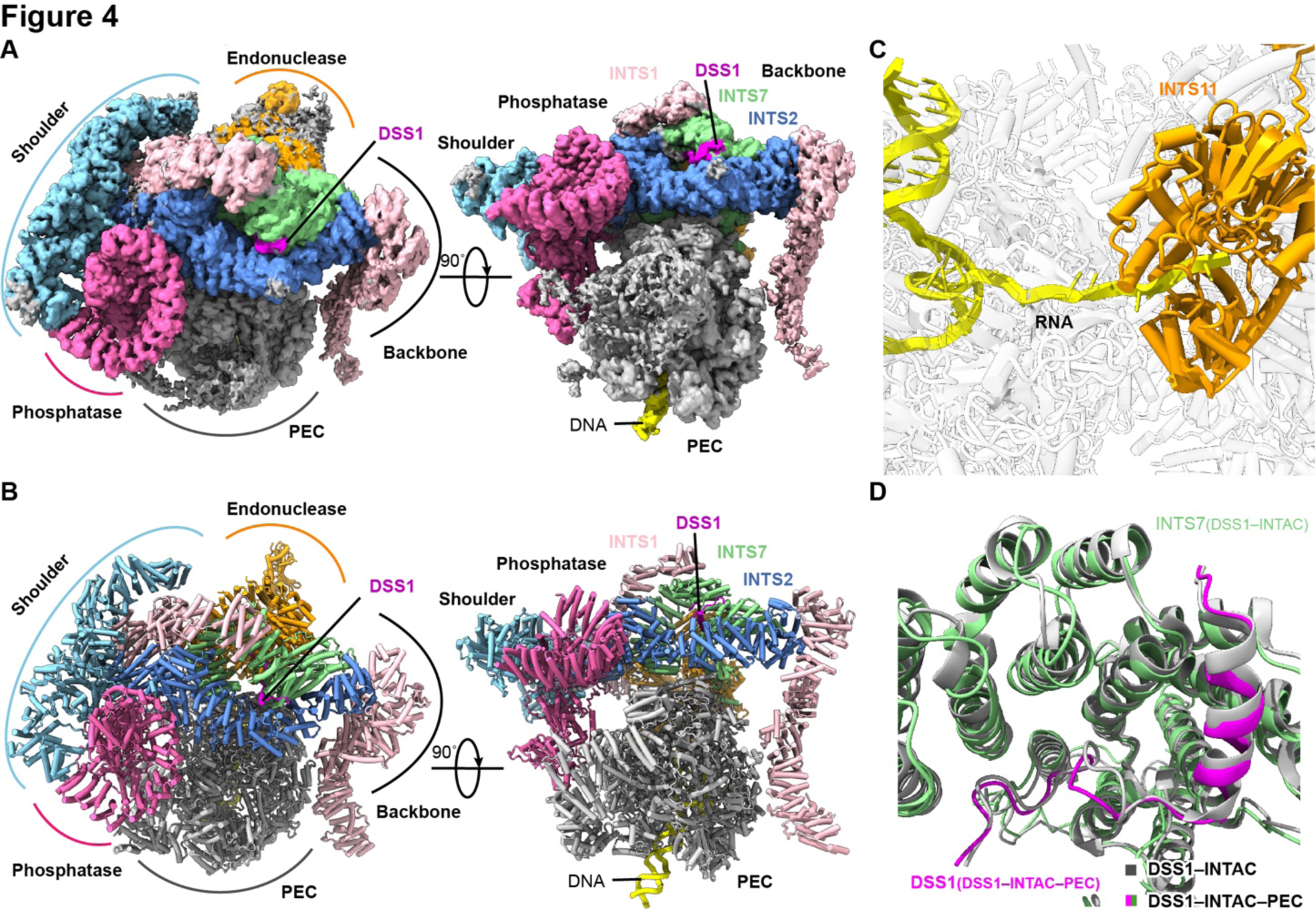
Structure of DSS1–INTAC–PEC. (A and B) Cryo-EM map (A) and molecular model (B) of DSS1-containing INTAC-PEC shown in two different views. Notably, the cryo-EM map is a composite map derived from multibody refinement. (C), Structural model showed close-up views of the mRNA (yellow) inserted into the catalytic center of INTS11 (orange). (D), Structural model showing the conformation of DSS1 in DSS1-INTAC (Gray) and DSS1-INTAC-PEC (INTS7: green, DSS1: magenta).

Examination of the DSS1–INTAC–PEC structure shows single-stranded RNA extending from the DNA–RNA hybrid within Pol II, and its 5’ end inserting into the catalytic pocket of INTS11, the enzymatic subunit of the INTAC endonuclease module (Fig. 4C). Overlaying INTS11 from the DSS1-containing INTAC and INTAC–PEC reveals a transition of the RNA endonuclease center from a closed, inactive state to an expanded, active conformation upon association with PEC (fig. S4D). Moreover, the phosphatase module exhibited structural reconfigurations within the INTAC–PEC framework (fig. S4, E and F), likely driven by the association of the phosphatase module, specifically INTS6, with NELF (*46, 47*). Notably, the conformation of DSS1, along with the backbone module, remains largely unaltered within INTAC and the INTAC–PEC assemblage (Fig. 4D). These findings support the notion that DSS1 is an integral subunit of INTAC, likely functioning as part of the INTAC backbone module.

### DSS1 is required for the stability of INTAC and its function in transcription

We next sought to determine whether DSS1 regulates the integrity of the INTAC complex by performing rescue experiments with DSS1-WT and DSS1-W39R, the latter of which affects DSS1’s interaction with INTAC but not the proteasome (fig. S3E). Specifically, we overexpressed either DSS1-WT or DSS1-W39R, followed by a time-course degradation of the endogenous DSS1. Western blotting of INTAC subunits showed a notable reduction in the protein levels of several examined subunits across different modules—backbone (INTS7), shoulder (INTS5 and INTS8), endonuclease (INTS4), and SOSS (INTS3)— upon introduction of DSS1-W39R compared to DSS1-WT in cells depleted of endogenous DSS1 (Fig. 5A). Using INTS3 as an example, we employed calibrated ChIP-seq with reference exogenous genome (ChIP-Rx) to estimate the chromatin occupancy of INTAC. Heatmap analysis showed a pervasive decline in INTS3 occupancy for all DSS1 bound peaks, including promoters (Fig. 5B and fig. S5A), an effect that can also be seen on example genes (Fig. 5C). These results demonstrate that DSS1 is required for the stability of INTAC.

**Fig. 5.**
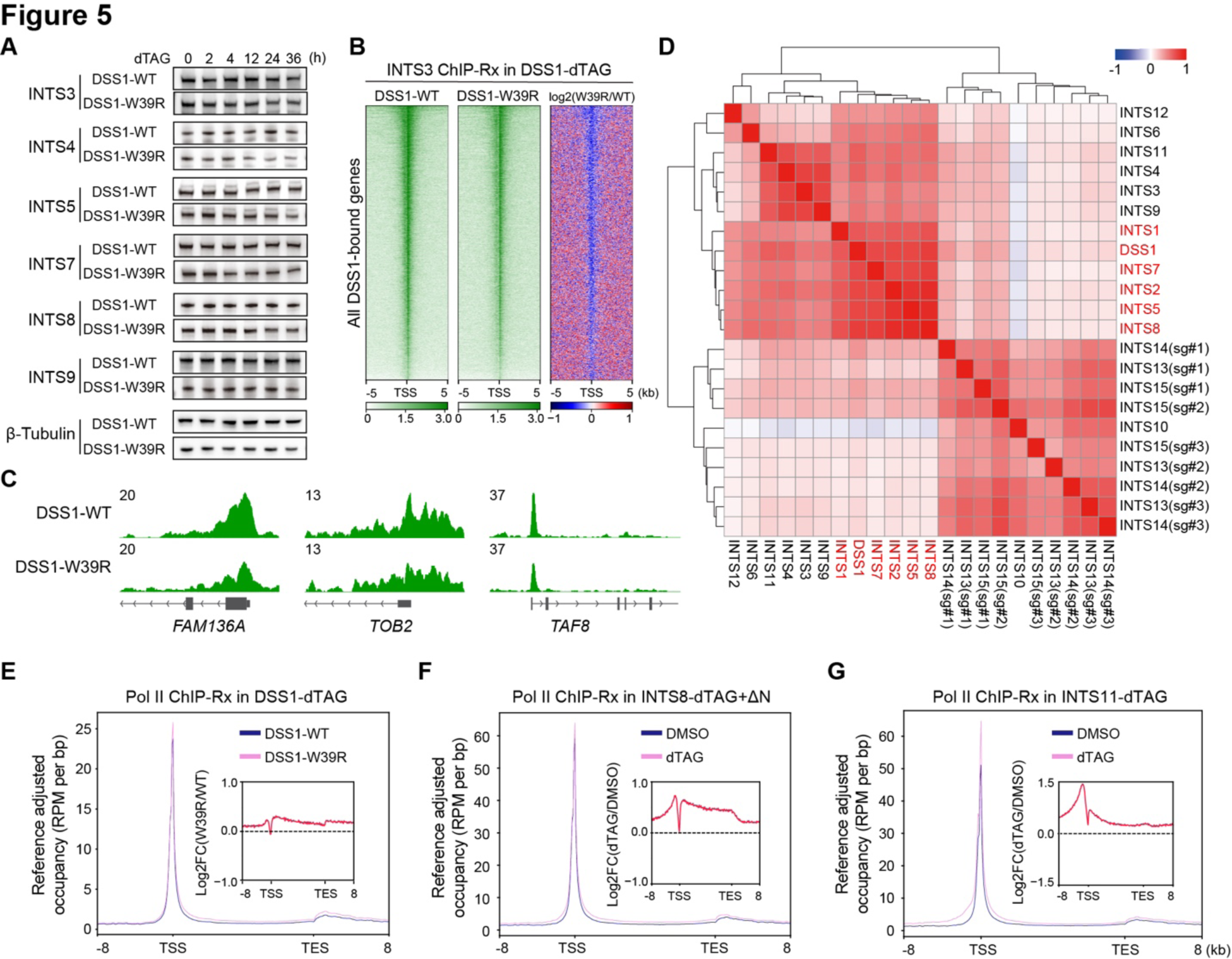
DSS1 regulates INTAC stability and its function in transcription. (A) Western blotting of INTAC subunits in DSS1-WT or DSS1-W39R expressing DSS1–Flag–dTAG cells with time-course dTAG treatment. (B) Heatmaps showing INTS3 ChIP-Rx signals over 10 kb regions centered on the TSS of target genes after endogenous DSS1 degradation in DSS1–Flag–dTAG cells with overexpression of DSS1-WT or DSS1-W39R. (C) Representative track examples of INTS3 occupancy in dTAG treated DSS1–Flag–dTAG cells with overexpression of DSS1-WT or DSS1-W39R. (D) Analysis of Perturb-seq showing the correlation of gene expression in cells lacking INTAC subunits. (E) Metagenes showing Pol ⅠⅠ ChIP-Rx signals in DSS1–Flag–dTAG cells overexpressing DSS1-WT or DSS1-W39R. (F and G) Metagenes showing Pol ⅠⅠ ChIP-Rx signals in INTS8-dTAG+ΔN (F) and INTS11-dTAG cells (G) treated with DMSO or dTAG.

To determine the role of DSS1 in supporting INTAC function in transcription, we analyzed genome-scale Perturb-seq results and found that depleting different modules of INTAC elicits distinct changes of gene expression (Fig. 5D) (*52*). Notably, the transcriptional changes following DSS1 depletion closely align with those seen in cells lacking subunits of the backbone and shoulder modules, which together play a scaffolding role in INTAC assembly (Fig. 5D).

We next performed Pol II ChIP-Rx in dTAG-treated DSS1–Flag–dTAG cells expressing either DSS1-WT or DSS1-W39R. Metagene and heatmap analysis showed increased Pol II occupancy across gene bodies in cells expressing the DSS1-W39R mutant compared to those with DSS1-WT (Fig. 5E and fig. S5B). We next analyzed the changes in Pol II occupancy in cells in response to rapid disruption of the INTAC shoulder and endonuclease modules (Fig. 5, F and G, and fig. S5, C and D) (*26*). This analysis shows that all three perturbations result in increased Pol II occupancy across gene bodies, with the alterations induced by DSS1 mutation being more similar to the changes seen in cells expressing the mutant INTS8, a scaffolding subunit of INTAC. These findings collectively underscore DSS1’s critical role in ensuring the stability and function of INTAC in transcriptional regulation.

## Discussion

In this study, we establish that human DSS1 is a *bona fide* component of the INTAC backbone module. Through our structural analysis of DSS1–INTAC, both alone and as part of a complex with the paused Pol II–DSIF–NELF elongation complex (PEC), we identify a close association between DSS1 and the INTS7 subunit of the INTAC backbone module. The detection of DSS1 in association with paused Pol II suggests its continuous engagement with INTAC throughout the premature termination process. A comparison of the DSS1–INTAC structures, with and without PEC, reveals that despite a reconfiguration in the overall structure of INTAC including activation of the RNA endonuclease module upon PEC interaction, DSS1’s positioning in relation to the backbone module remains unchanged in the presence of PEC, thus underscoring a strict scaffolding role for DSS1.

Importantly, we identified a specific point mutation in DSS1 that disrupts its interaction with INTAC, without affecting its association with the 26S proteasome, and this point mutant significantly impairs the chromatin enrichment of DSS1. This suggests that chromatin-bound DSS1 primarily exists within INTAC. Expression of this mutant destabilizes multiple INTAC subunits across different modules, further demonstrating the importance of DSS1 in promoting the complex’s stability. Moreover, disrupting the DSS1–INTAC interaction triggers transcriptional alterations akin to those seen in INTAC deficient cells, affirming DSS1’s critical function in facilitating INTAC’s role in transcriptional regulation.

Among the four established DSS1-associated complexes—26S proteasome, TREX-2, BRCA2, and INTAC—the exact role of DSS1, whether acting independently within each complex or contributing to a shared biological outcome, remains an open question. The possibility that the regulation of DSS1 levels could enable a coordinated cellular response across these complexes is an intriguing hypothesis. Supporting this idea, each of these complexes plays a role, either directly or indirectly, in genome stability. For example, BRCA2 plays a critical role in DSB repair through homologous recombination and helps mitigate the accumulation of R-loops, which can induce replication stress and genomic instability due to the susceptibility of ssDNA to damage and the obstruction of replication fork progress by DNA:RNA hybrids. BRCA2’s involvement in R-loop mitigation is linked to its interaction with the TREX-2 subunit PCID2, with both being direct partners of DSS1 (*53–55*).

While it is improbable for a single DSS1 molecule to simultaneously engage with BRCA2 and TREX-2 due to the spatial constraints of their binding sites, the potential role of DSS1 in bridging BRCA2 and TREX-2’s activities against R-loops has yet to be clarified. Similarly, INTAC was recently found to recognize and attenuate R-loops that arise at sites of paused Pol II, thus preventing R-loop-related genomic instability (*27, 56*). Moreover, evidence suggests that DSS1 is recruited along with the 26S proteasome to DSBs to facilitate their repair (*57, 58*). These observations suggest that DSS1’s involvement across these complexes might reflect multifaceted yet interconnected biological functions. Exploring how these complexes might collaborate, especially through the regulation of DSS1’s availability or stability, presents an intriguing area for future research.

## Materials and Methods

### Reagents, materials and cell culture

Detailed information for reagents and materials, including antibodies and cell lines, used in this study is provided in Table S2.

Human DLD-1 cells were cultured in McCoy’s 5A medium (BasalMedia) supplemented with 10% fetal bovine serum (FBS; Yeasen) and 1× penicillin-streptomycin (Meilunbio). HEK293T cells were maintained in Dulbecco’s modified Eagle medium (DMEM, BasalMedia) supplemented with 10% FBS and 1× penicillin-streptomycin.

HEK 293 Flp-In T-Rex cells were cultured in high glucose (4.5 mg/ml) DMEM medium containing 10% FBS and 1× penicillin/streptomycin. Additionally, 100 μg/ml zeocin and 15 μg/ml blasticidin were supplied in DMEM medium to maintain wild-type 293 Flp-In T-Rex cells cells.

### Molecular cloning

The human DSS1 gene was amplified from a cDNA library reverse transcribed from SK-HEP1 cells. The DSS1 gene was subcloned into a modified pcDNA5/FRT/TO plasmid (Invitrogen), resulting in an N-terminal 2x Strep and 3x FLAG tag, yielding the pcDNA5-StFLAG-DSS1 plasmid. To produce lentivirus for gene overexpression assays, the DSS1 gene and it mutant (W36R) were also subcloned in to pLVX plasmid, resulting in pLVX-DSS1-WT or –Mutant.

### Genome editing for dTAG endogenous knock-in

The procedures for generating the dTAG endogenous knock-in cell line were conducted following established criteria (*59, 60*). DLD-1 cell suspension (1×10^6^ cells) was combined with PITCh plasmids, including PX459-sgRNA for targeting specific genomic sites within target genes, pCRISPR-PITChv2 containing the dTAG microhomology repair template, and sg-PITCh for cutting and fetching out the repair template. Electroporation was then employed, followed by a 2-day recovery period in regular culture medium. After the recovery phase, cells underwent gradient dilution and were cultured with 1–2 μg/ml puromycin (Meilunbio) for 10–14 days. Surviving single colonies were selected, expanded, and subjected to genotype confirmation. The effectiveness of protein degradation in positive clones was validated through dTAG treatment, followed by western blots analysis.

### Generation of stable overexpression cell lines

To produce lentivirus for gene overexpression assays, HEK293T cells underwent co-transfection with plasmids pLVX-DSS1-WT & Mutant (or the empty pLVX vector plasmid as the control), which harbored the blasticidin resistance gene, along with psPAX2 and pMD2.G. The transfection plasmids were mixed in a 4:2:1 ratio in Opti-MEM medium utilizing the polycationic agent polyethylenimine (PEI; Sigma-Aldrich). Following transfection, the HEK293T culture supernatant, containing viral particles, was harvested at 48 h and 72 h, subsequently filtered through a 0.45 μm filter.

DLD-1 Cell infection with lentivirus was conducted in the presence of 10 mg/ml Polybrene (YEASEN) for 24 h. Infected cells underwent treatment with 10 μg/ml blasticidin for 7–10 days before collection. Evaluation of overexpression efficiency was performed through western blots.

### Co-immunoprecipitation assays

For co-immunoprecipitation (Co-IP) assays, DLD-1 cells (1–2×10^7^) were harvested by scraping and subsequently washed twice with ice-cold phosphate-buffered saline (PBS). In experiments involving the comparison of multiple cell lines, normalization by weight was implemented to block variations in cellular mass. The cell pellet was resuspended in 900 μl of ice-cold lysis buffer (20 mM Tris-HCl, pH 8.0, 150 mM NaCl, 1 mM EDTA, 0.2– 0.5% NP40, 10% glycerol, 1× protease inhibitor) and gently rotated at 4 ℃ for 1 h.

The resulting lysate was clarified by centrifugation at 20,000 g for 20 min at 4 ℃. The supernatant was then incubated with 2–5 μg of the relevant antibody for each immunoprecipitation reaction, followed by 9.5 h of rotation at 4 ℃. rProtein A/G MagPoly Beads (Smart-Lifesciences), pre-blocked with 1 mg/ml BSA for 1 h, were introduced to the samples, and the mixture was rotated for an additional 3 hours at 4 ℃. Following incubation, samples with beads underwent collection using a magnetic rack, and the beads were subjected to four washes with lysis buffer. Finally, the samples were eluted by adding 50–100 μl of 1× SDS loading buffer, followed by subsequent western blot analysis.

### Immunofluorescence analysis

DLD-1 cells were cultured on coverslips for a minimum of 24 h preceding the experimental procedures. Following a wash with phosphate-buffered saline (PBS), cells underwent fixation with 4% paraformaldehyde (PFA) for a duration of 10 minutes. After three consecutive PBS washes, cell permeabilization was achieved by treatment with 0.5% Triton X-100 in PBS for 10 min, followed by blocking with 4% bovine serum albumin (BSA) in PBS for 30 min.

Primary antibodies, appropriately diluted in ice-cold 4% BSA according to the recommended ratios by the manufacturers, were applied to the cells for an overnight incubation at 4 ℃. Subsequent to three PBS washes, cells underwent a 1-hour incubation with the respective secondary antibodies. Following this, cells were mounted in ProLong Gold Antifade Mountant with DAPI (Invitrogen) before imaging.

Image acquisition was performed using the Leica TCS SP8 laser-scanning confocal microscopy. Unless explicitly stated otherwise, all procedures were executed at room temperature. The division of nuclei and cytoplasmic regions, as well as the quantification of immunofluorescence intensity, were executed using the Fiji built on ImageJ2 2.14.0/1.54f, complemented by statistical analysis conducted through GraphPad 9.5.1 Prism.

### Western blotting

The samples underwent lysis using 1× SDS loading buffer, followed by heating at 99 ℃ for 10 minutes. Subsequently, the lysates were loaded onto a FuturePAGE™ 4–20% 15 Wells SDS page (ACE Biotechnology) and transferred to an Immobilon®-NC Transfer Membrane (Millipore). Blots were then blocked using Protein Free Rapid Blocking Buffer (Epizyme Biotech) for 20 minutes. After that they were subjected to overnight incubation at 4 ℃ with appropriately diluted primary antibodies. Following three washes with TBS containing Tween-20 (TBST), blots were incubated for 1 hour at room temperature with HRP-conjugated anti-rabbit– or anti-mouse-IgG (Abclonal), diluted at 1:5000 in TBST, followed by three additional washes in TBST.

Signal detection was achieved using Immobilon Western Chemiluminescent HRP Substrate (Millipore), and images were captured using a Tanon Imaging system. Subsequent quantification of band intensity was performed using Fiji built on ImageJ2 2.14.0/1.54f.

### ChIP-Rx

ChIP-Rx was performed following established protocols (*61*). Each immunoprecipitation (IP) utilized 1×10^7^ cells, which were crosslinked with 1% formaldehyde (Sigma) for 10 minutes at room temperature. The crosslinking reaction was terminated by the addition of glycine (final concentration 125 mM) and subsequent incubation for 5 minutes. Cells were then rinsed twice with ice-cold PBS, scraped, and collected by centrifugation.

The cell pellet was resuspended in Lysis buffer (50 mM HEPES pH 7.4, 150 mM NaCl, 2 mM EDTA, 0.1% SDS, 0.1% sodium deoxycholate, 1x protease inhibitor (Roche)) and subjected to sonication (Qsonica) to fragment chromatin to an appropriate size range. The soluble fraction was collected by centrifugation and mixed with 25% of lysate from mouse embryonic stem cells (mESCs) processed identically as a spike-in for normalization.

The mixed lysate was then incubated with antibodies overnight and BSA-blocked magnetic Protein A/G beads for 2 h, followed by three washes with High-salt Wash buffer (20 mM HEPES pH 7.4, 500 mM NaCl, 1 mM EDTA, 1.0% NP-40, 0.25% sodium deoxycholate), two washes with Low-salt Wash buffer (20 mM HEPES pH 7.4, 150 mM NaCl, 1 mM EDTA, 0.5% NP-40, 0.1% sodium deoxycholate), and a final wash with TE buffer containing 50 mM NaCl.

Beads were then eluted with Elution buffer (50 mM Tris-HCl pH 8.0, 10 mM EDTA, 1.0% SDS). The eluted samples were treated with Protease K, and DNA was purified using the Phenol/Chloroform/Isoamyl Alcohol extraction method. Subsequently, libraries for sequencing were prepared using the VAHTS Universal Plus DNA Library Prep Kit for Illumina (Vazymes). The libraries were sequenced using NovaSeq 6000 platform (Mingma Technologies).

### CUT&Tag

The CUT&Tag procedure was optimized in accordance with a previously published protocol (*62*). DLD-1 cells were harvested using Accutase (Thermo Fisher Scientific) to prevent overdigestion. For each CUT&Tag reaction, 5 × 10^5^ cells were typically utilized to ensure sufficient DNA yield for subsequent library construction. Following centrifugation (600 g, 3 minutes) at room temperature, cells were washed twice with 800 μl of wash buffer (20 mM HEPES pH 7.5, 150 mM NaCl, 0.5 mM spermidine, 1× protease inhibitor) and subsequently resuspended in 100 μl of wash buffer within low-retention PCR tubes.

The concanavalin-A-coated magnetic beads (Smart-Lifesciences) were pre-activated and resuspended in an equal volume of binding buffer (20 mM HEPES pH 7.5, 10 mM KCl, 1 mM CaCl_2_, 1 mM MnCl_2_). Subsequently, 10 μl of activated concanavalin A beads were added to 5 × 10^5^ cells, followed by a 10-minute incubation period under gentle rotation. The bead-bound cells were magnetized to remove excess liquid, then resuspended in 50 μl of antibody buffer (20 mM HEPES pH 7.5, 150 mM NaCl, 0.5 mM spermidine, 1× protease inhibitor, 0.05% digitonin, 0.01% NP-40, 2 mM EDTA).

Next, 1 μg of either Flag (Abclonal) or HA (proteintech) antibody was added to facilitate DSS1 binding, and the mixture was rotated overnight at 4 ℃. For IgG control experiments, mouse or rabbit IgG was substituted accordingly. Following overnight incubation, the bead-bound cells were subjected to successive incubations with rabbit anti-mouse IgG (Solarbio, diluted 1:100) and mouse anti-rabbit IgG (Solarbio, diluted 1:100) in 100 μl of antibody buffer for 1 hour at room temperature. Subsequently, the cells were washed four times with dig-wash buffer (antibody buffer without 2 mM EDTA) to remove unbound antibodies.

The pAG-Tn5 adapter complex was prepared in dig-300 buffer (20 mM HEPES pH 7.5, 300 mM NaCl, 0.5 mM spermidine, 1× protease inhibitors, 0.01% digitonin, 0.01% NP-40) to a final concentration of 0.2 μM. The bead-bound cells were resuspended in 100 μl of pAG-Tn5 mix and incubated at room temperature for 1 hour before supernatant removal. Following thorough washing, the tagmentation reaction was initiated in 40 μl of tagmentation buffer (10 mM TAPS-KOH pH 8.3, 10 mM MgCl2, 1% DMF) at 37 ℃ for 1 h.

To halt the reaction, 1.5 μl of 0.5 M EDTA, 0.5 μl of 10% SDS, and 1 μl of 20 mg/ml protease K were added. After a 1-hour incubation at 55 ℃, DNA purification was performed using VAHTS DNA Clean Beads (Vazyme), with elution in 10 μl of 0.1% Tween-20. The eluent was subsequently mixed with 10 U of Bst 2.0 WarmStart DNA polymerase (NEB) and 1× Q5 polymerase reaction buffer (NEB) in a 20 μl reaction system.

The reaction was carried out at 65 ℃ for 30 min, followed by an additional 20 min at 80 ℃ to inactivate the Bst 2.0 WarmStart DNA polymerase. The purified DNA was then subjected to amplification using Q5 high-fidelity DNA polymerase (NEB) with a universal i5 primer and a uniquely barcoded i7 primer. The exact number of PCR cycles was determined by qPCR prior to amplification, with 11–12 cycles typically sufficient to yield an adequate library for sequencing.

Following library size-selection with 0.56–0.85 VAHTS DNA Clean Beads to achieve library sizes ranging from 200 to 700 bp, the products were sequenced on the NovaSeq 6000 platform (Mingma Technologies).

### Native DSS1 sample purification and LC-MS

To establish stable DSS1 overexpression HEK 293/Flp-In/T-Rex cell lines, cells were pre-cultured in a 10-cm dish for one day. When reaching 50% confluency, cells were co-transfected with 0.5 μg pcDNA5-StFLAG-DSS1 and 4.5 μg recombinase plasmid pOG44 (Invitrogen) for 48 hours. Subsequently, cells were passaged and subjected to a 14-day selection with 200 μg/ml hygromycin B. Western blot (WB) was employed to validate the resulting cell lines.

The purification process was adapted from previous studies (*63, 64*). In brief, a total of 50 dishes (15 cm) of DSS1-overexpressed HEK 293/Flp-In/T-Rex cells were used for each purification. When the cells reached 70% confluency, DSS1 expression was induced with 1 μg/ml tetracycline for 24 hours. Cells was collected with a cell scraper, washed twice with cold 1× PBS buffer, and resuspended in lysis buffer (20 mM HEPES pH 7.4, 100 mM KOAc, 5 mM MgCl_2_, 1 mM DTT, 0.5 mM NaF, 1 mM Na_3_V_3_O_4_, 0.4% NP–40, 1× protease inhibitor mix). Cells were lysed by 10–15 strokes using a 15 ml douncer. The cell lysate was clarified by centrifugation at 10,000 × g at 4 ℃ for 15 minutes, and the supernatant was incubated with 200 μl Flag beads (A2220, Sigma) in a 50 ml centrifuge tube for 2 hours. The beads were then transferred to a small chromatography column (Cytiva, 27-5140-01), washed once with lysis buffer and three times with wash buffer (20 mM HEPES pH 7.4, 100 mM KCl, 5 mM MgCl_2_, 0.04% NP-40). Finally, the protein complex was eluted by incubating with 500μl elution buffer (20 mM HEPES pH 7.4, 100 mM KCl, 5 mM MgCl_2_, 0.4 mg/ml 3×Flag peptide) for 45 minutes.

The eluate was concentrated using a 100 kDa cut-off concentrator (Millipore). The purified sample was checked by SDS-PAGE gel electrophoresis and Coomassie Brilliant Blue staining. Subsequently, protein quantification and identification were performed through label-free protein quantification using LC-MS.

### Protein expression and purification

Protein expression and purification were performed essentially as previously described (*65*). In brief, the 14 full-length open reading frames (ORFs) of human INTAC subunits (INTS1 to INTS14) were separately cloned into a modified pCAG vector. INTS2, INTS3, INTS4, and INTS10 were tagged with an N-terminal Protein A (ProA) tag. To purify DSS1 containing INTAC complex, pLVX-DSS1-WT plasmid was co-expressed together with all 14 INTAC subunits plasmids in Expi293F cells and purified by immunoglobulin G (IgG) affinity chromatography and on-column digestion. The eluate was concentrated for further purification by glycerol density gradient sedimentation. The fractions, 200 μl each, were collected manually from the top of the gradient and analyzed by SDS-PAGE. Peak fractions were pooled and concentrated to 1 to 2 mg/ml accompanied with the removal of glycerol. The purified complex was used for cryo-EM sample preparation. Pol II was isolated from *S. scrofa* thymus and purified following the reported protocol (*66*). Four residue substitutions (G882S of RPB2, T75I of RPB3, S140N of RPB3, and S126T of RPB6) exist between *S. scrofa* and *H. sapiens* Pol II. Human NELF and DSIF were separately expressed and purified as previously described (*46*).

### EM sample preparation

To prepare the DSS1–INTAC complex for cryo-EM, the purified DSS1–INTAC sample was further cross-linked and purified using gradient fixation (Grafix) as described (*65, 67*). The fractions were collected manually and the homogeneity of peak fractions was assessed by negative-stain EM. Fractions of interest were concentrated, followed by cryo-EM grid preparation.

The DSS1–INTAC–PEC complex for cryo-EM was prepared essentially as previously described (*46*). In brief, the purified *S. scrofa* Pol II (275 pmol) and the RNA-template DNA hybrid (550 pmol) were incubated for 15 min at 30 ℃, shaking at 300 rpm, followed by the addition of non-template DNA with a twofold molar excess relative to template DNA and further incubation for 15 min at 30 ℃. The purified DSIF and NELF were added in a twofold molar excess relative to Pol II for the PEC reconstitution. The sample was incubated for 1 hour at 4 ℃, followed by the addition of the purified DSS1–INTAC (230 pmol) and incubation for another 2 hours at 4 ℃. The resulting sample was subjected to GraFix (*67*). The fractions were collected manually and the homogeneity of peak fractions was assessed by negative-stain EM. Finally, fractions of interest were concentrated, followed by cryo-EM grid preparation.

For negative-stain EM, 5 μl of freshly purified protein sample was applied onto a glow-discharged copper grid supported by a thin layer of carbon film for 1 min before negative staining by 2% (w/v) uranyl formate at room temperature. The negatively stained grid was loaded onto a FEI Talos L120C microscope operated at 120 kV, equipped with a Ceta CCD camera.

For cryo-EM grid preparation, 4 μl of protein sample (about 0.7 mg/ml for DSS1–INTAC complex, about 0.9 mg/ml for DSS1–INTAC–PEC complex) was applied onto a glow-discharged holey carbon grid (Quantifoil Au, R1.2/1.3 or R2/2, 300 mesh). After blotting for 3 to 6 s, the grid was vitrified by plunging it into liquid ethane using a VitrobotMark IV (FEI) operated at 4 ℃ and 100% humidity.

### Cryo-EM data collection and image processing

Cryo-EM data were acquired using a Titan Krios electron microscope (FEI) operating at 300kV at the Cryo-EM platform of Peking University. The microscope was equipped with a K2 direct detector (Gatan) and a GIF quantum energy filter (Gatan) set to a slit width of 20 eV. Automated data acquisition was performed using Serial EM software in super-resolution mode at a nominal magnification of 130,000×, corresponding to a calibrated pixel size of 1.055 Å and 1.054 Å for the DSS1–INTAC and DSS1–INTAC–PEC sample, respectively. The defocus range was set from –1.5 to –2.5 μm. Each image stack was dose fractionated into 32 frames with a total exposure dose of about 50 e−/Å^2 and an exposure time of 6.96 s and 6.72 s, respectively.

All the image stacks were motion-corrected and dose-weighted using MotionCorr2 (*68*). The contrast transfer function (CTF) parameters were estimated by GCTF (*69*) from non-dose weighted micrographs. Micrographs with an estimated resolution of less than 5 Å and astigmatism of less than 5% were manually inspected for contamination or carbon breakage.

For the DSS1–INTAC dataset, 587,397 particles were picked from 3,965 good micrographs using Gautomatch (https://www2.mrc-lmb.cam.ac.uk/download/gautomatch-053/) with the published INTAC structure (PDB: 7CUN) (*70*) as a reference. These particles were extracted in Relion (*71*) and then imported into cryoSPARC (*72*).

Heterogeneous refinement in cryoSPARC (*72*) using the published INTAC structure as references (*70*) yielded 95,815 particles representing the INTAC structure. After 3D classification in Relion (*71*), 43,790 particles showing the intact INTAC structure (termed DSS1–INTAC) were identified. Final 3D refinement and CTF refinement resulted in a reconstruction at 4.1 Å resolution, based on the gold-standard Fourier shell correlation (FSC) 0.143 criterion. To improve local resolution, the INTAC map was divided into three bodies: the phosphatase and shoulder part, the endonuclease module, and the backbone of the INTAC, followed by multibody refinement in Relion (*71*). The detailed sorting scheme is shown in Fig. S1.

For the DSS1–INTAC–PEC dataset, an identical procedure was followed. Initially, 843,523 particles were picked from 7,585 good micrographs using Gautomatch (https://www2.mrc-lmb.cam.ac.uk/download/gautomatch-053/) with the published INTAC–PEC structure (PDB: 7YCX) (*46*) as a reference. After processing in cryoSPARC (*72*) and Relion (*71*), 26,822 particles showing the INTAC associated with the paused elongation complex (DSS1–INTAC–PEC) were identified. Final 3D refinement and CTF refinement yielded a reconstruction at 4.6 Å resolution. Due to the high flexibility of the paused elongation complex (PEC), the DSS1–INTAC–PEC structure was divided into four bodies: the phosphatase and shoulder part, the endonuclease module, the backbone module, and the PEC, followed by multibody refinement in Relion (*71*). The data processing steps are summarized in Fig. S1.

All the visualization and evaluation of 3D density maps were performed with UCSF Chimera (*73*) or UCSF ChimeraX (*74*). All final maps were post-processed and local resolution filtered using masks automatically generated in Relion (*71*). Simultaneously, the final maps were sharpened using DeepEMhancer (*75*).

### Model building and structure refinement

The structural models of DSS1–INTAC and DSS1–INTAC–PEC were built based on their cryo-EM maps and corresponding focused refined maps. Human INTAC (PDB: 7CUN) (*70*) and INTAC–PEC (PDB: 7YCX) (*46*) structures were initially docked into the cryo-EM maps of DSS1–INTAC and DSS1–INTAC–PEC using rigid body fitting in UCSF Chimera (*73*), followed by manual adjustments in COOT (*76*). The INTAC and PEC components in both models remained unchanged, while the DSS1 component was manually built in COOT (*76*) according to its corresponding density maps.

Due to the low overall resolution, the rigid body fitted model for DSS1–INTAC–PEC was used as the final model without any further refinement. However, the structural model of the DSS1–INTAC complex was refined against the 4.1 Å overall map in real space with PHENIX (*77*) and was subsequently validated using MolProbity (*78*). The statistics of the map reconstruction and model refinement are summarized in Table S1. Each focused refined maps from multibody refinement were used to create the composite map using PHENIX (*77*), which was then used to create the figures using UCSF ChimeraX (*74*).

### CUT&Tag data analysis

Raw CUT&Tag reads were trimmed using Trim Galore v.0.6.6 (Babraham Institute). Trimmed reads were aligned to the human hg19 genome using Bowtie v.2.4.4 with the following parameters: “-N 1 –L 25 –X 700 –-no-mixed –-no-discordant” (*79*). Duplicate reads were removed with Picard Tools v2.26.6 (Broad Institute). The reads were then shifted to compensate for the offset in tagmentation site relative to the Tn5 binding site using the alignmentSieve function of deepTools v.3.5.1 with the ‘--ATACshift’ option (*80*). Peak calling was performed by macs2 v2.2.6 with a q-value threshold of 0.05 (*81*).

### ChIP-Rx data analysis

The raw ChIP-Rx reads were trimmed by Trim Galore v0.6.6 (Babraham Institute) and aligned to the human hg19 and mouse mm10 assemblies using Bowtie v2.3.5.1 with default parameters (*79*). Low mapping quality reads (MAPQ < 30) and PCR duplicates were removed using SAMtools v1.9 (*82*) and Picard v2.26.6 (Babraham Institute). The read number for each of the ChIP-Rx sample were then collected using SAMtools v1.9 (*82*), and the normalization factor was calculated as 1e6/spike-in_count. Normalized bigwig files were generated by deeptools v3.5.0 (*80*).

The Materials and Methods section should provide sufficient information to allow replication of the results. Begin with a section titled Experimental Design describing the objectives and design of the study as well as prespecified components.

## Author contributions

Conceptualization: CFX, CJ, Xu Y, XC. Investigation: XC, ZQX, ZH, SA, ZWY, Xiong Y, HZ. Visualization: XC, ZQX, ZH, SA, CJ, CFX. Supervision: CFX, XC. Writing: CFX, CJ, Xu Y, XC

## Data and materials availability

Sequencing data will be deposited upon acceptance of the manuscript. Raw sequencing data have been deposited at GEO with accession number of GSEXXXXXX. The cryo-EM maps and coordinates have been deposited to the Electron Microscopy Data Bank (EMDB) and Protein Data Bank (PDB), including DSS1– INTAC (EMD-XXXXX, PDB ID XXXX) and DSS1–INTAC–PEC (EMD-XXXXX). All data needed to evaluate the conclusions in this paper are present in the paper, the Supplementary Materials, and deposited GEO dataset and structures.

