## Supplemental figures and table 1 for "Stabilization of Integrator/INTAC by the small but versatile DSS1 protein"

Figure S1

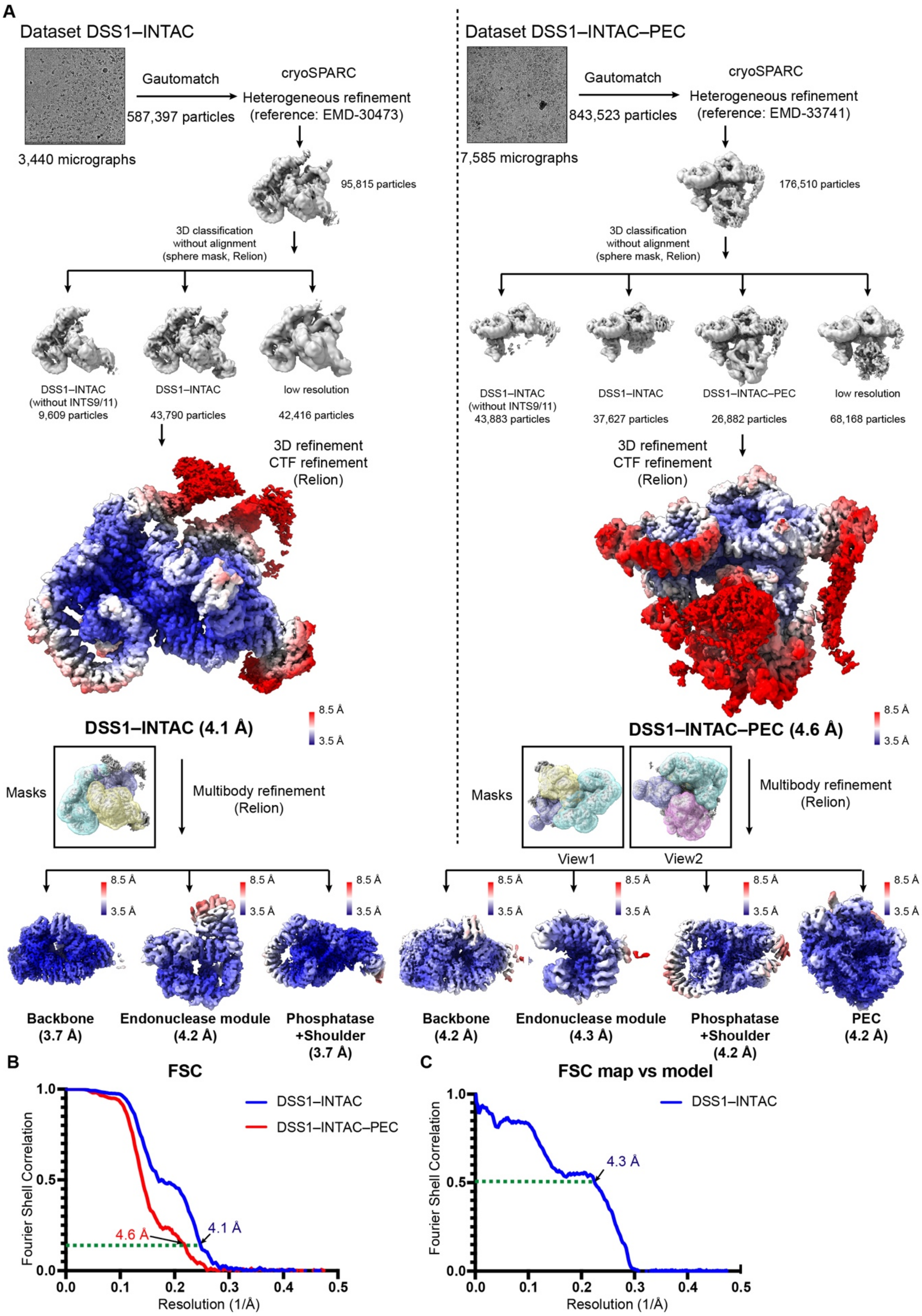
A
Dataset DSS1–INTAC
Gautomatch
587,397 particles
3,440 micrographs
cryoSPARC
Heterogeneous refinement
(reference: EMD-30473)
95,815 particles
3D classification
without alignment
(sphere mask, Relion)
DSS1–INTAC
(without INTS9/11)
9,609 particles
DSS1–INTAC
43,790 particles
low resolution
42,416 particles
3D refinement
CTF refinement
(Relion)
DSS1–INTAC (4.1 Å)
8.5 Å
3.5 Å
Masks
Multibody refinement
(Relion)
Backbone
(3.7 Å)
Endonuclease module
(4.2 Å)
Phosphatase
+Shoulder
(3.7 Å)
Dataset DSS1–INTAC–PEC
Gautomatch
843,523 particles
7,585 micrographs
cryoSPARC
Heterogeneous refinement
(reference: EMD-33741)
176,510 particles
3D classification
without alignment
(sphere mask, Relion)
DSS1–INTAC
(without INTS9/11)
43,883 particles
DSS1–INTAC
37,627 particles
DSS1–INTAC–PEC
26,882 particles
low resolution
68,168 particles
3D refinement
CTF refinement
(Relion)
DSS1–INTAC–PEC (4.6 Å)
Masks
View1
View2
Multibody refinement
(Relion)
Backbone
(4.2 Å)
Endonuclease module
(4.3 Å)
Phosphatase
+Shoulder
(4.2 Å)
PEC
(4.2 Å)
B
FSC
DSS1–INTAC
DSS1–INTAC–PEC
4.6 Å
4.1 Å
Fourier Shell Correlation
Resolution (1/Å)
C
FSC map vs model
DSS1–INTAC
4.3 Å
Fourier Shell Correlation
Resolution (1/Å)

**Fig. S1. Cryo-EM structure determination of DSS1–INTAC and DSS1–INTAC–PEC.**

(A) The sorting schemes of the cryo-EM datasets for the DSS1–INTAC sample (left of the dash line) and DSS1–INTAC–PEC sample (right of the dash line). The masks and software used during data processing are shown on the side. The final DSS1–INTAC and DSS1–INTAC–PEC maps are colored according to their local resolution distribution. (B and C) The FSC curves (B) of the maps, along with the model-to-map correlation curve (C, applicable only to DSS1–INTAC), were calculated using Relion. The resolution was estimated using either the 0.143 or 0.5 cutoff criterion (indicated by the green line).

**Figure S2**

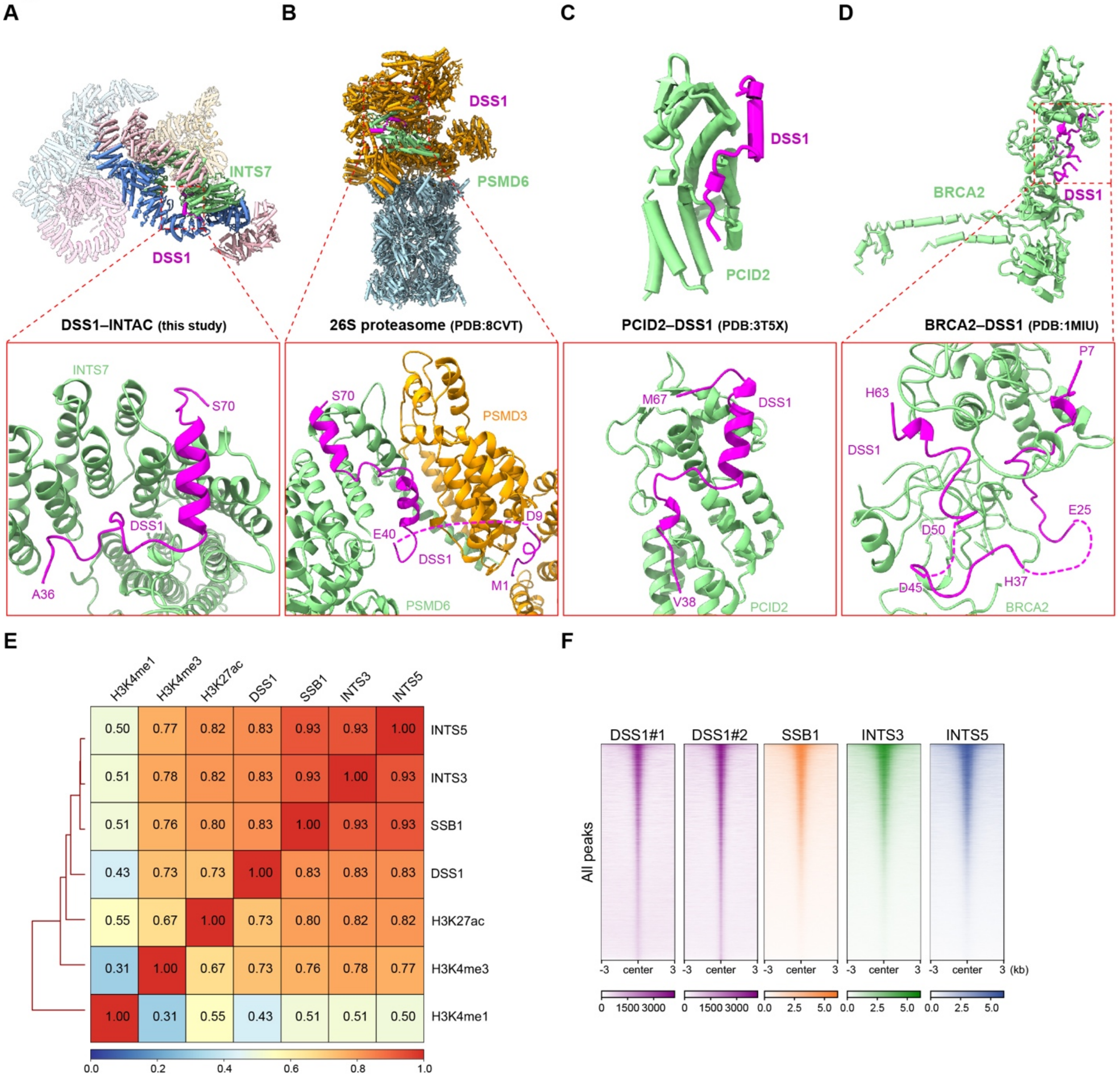

**Fig. S2. Structural comparison of DSS1-containing complexes and correlation analysis of INTAC subunits including DSS1.**

(A to D) Overall (top) and close-up views (bottom) of structures of DSS1-containing complexes, including DSS1–INTAC (A), 26S proteasome (B), TREX-2 components PCID2–DSS1 (C), and BRCA2–DSS1 (D). The internal missing regions of the DSS1 are connected by dash lines. (E) Correlation analysis for the genomic occupancy of DSS1, SSB1, INTS3, INTS5, H3K27ac, H3K4me3, and H3K4me1. The numbers indicate Pearson correlation coefficients. (F) Occupancy of DSS1, SSB1, INTS3, INTS5 over 6 kb regions centered on all peaks of DSS1.

### Figure S3

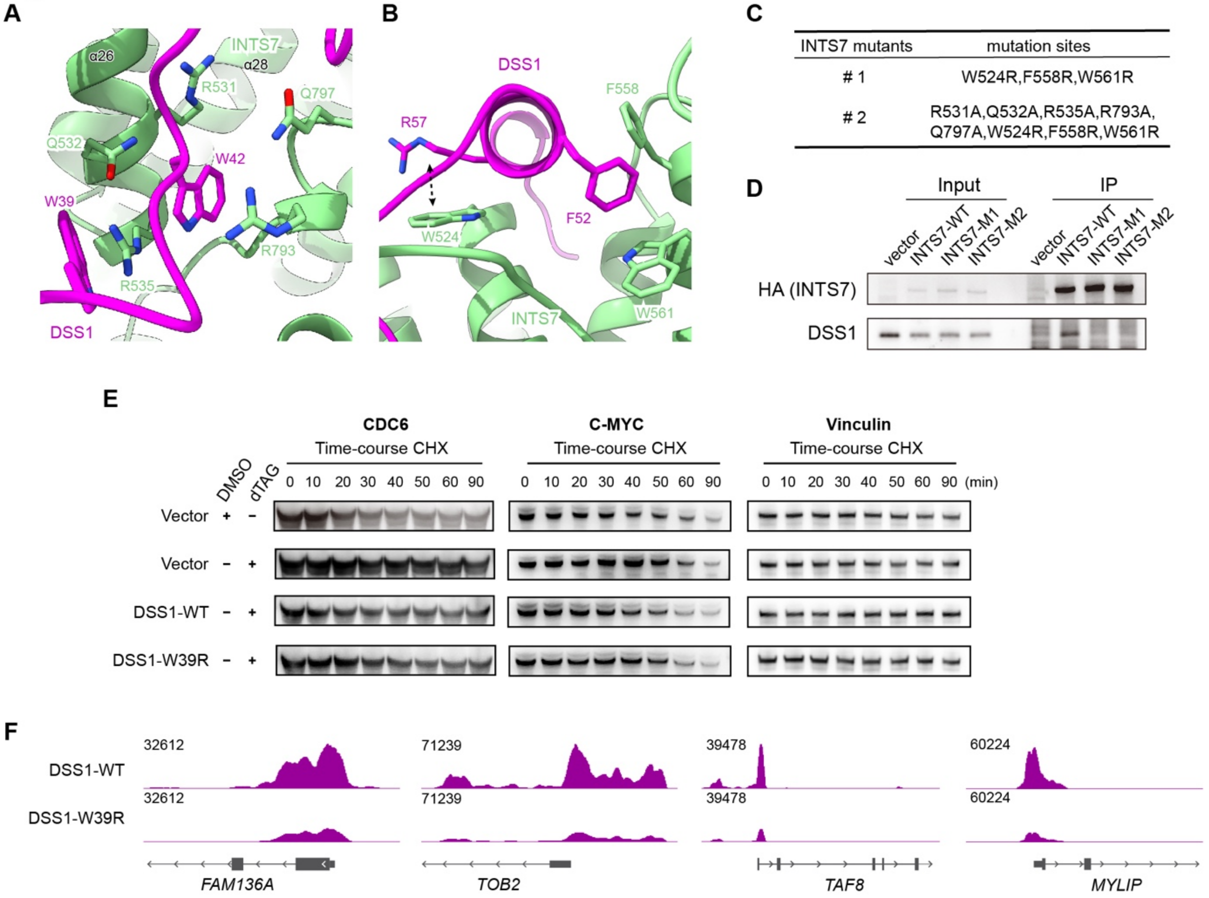

**Fig. S3. Separation of DSS1's implication in INTAC and proteasome.**

(A and B) Close-up views presents a detailed interactions between DSS1 (magenta) and INTS7 (green) at two different interfaces. (C) Mutating essential residues of INTS7 for its interaction with DSS1 highlighted in (A) and (B). (D) Co-IP analysis of HA (INTS7) in cells overexpressing wild-type INTS7 (INTS7-WT), or the two INTS7 mutants shown in (C). (E) Time-course cycloheximide (CHX) administration in the dTAG treated DSS1–Flag–dTAG cells with the overexpression of DSS1-WT, DSS1-W39R, or vector. (F) Representative track examples showing the occupancy of DSS1 in dTAG-treated DSS1–Flag–dTAG cells expressing DSS1-WT or DSS1-W39R.

### Figure S4

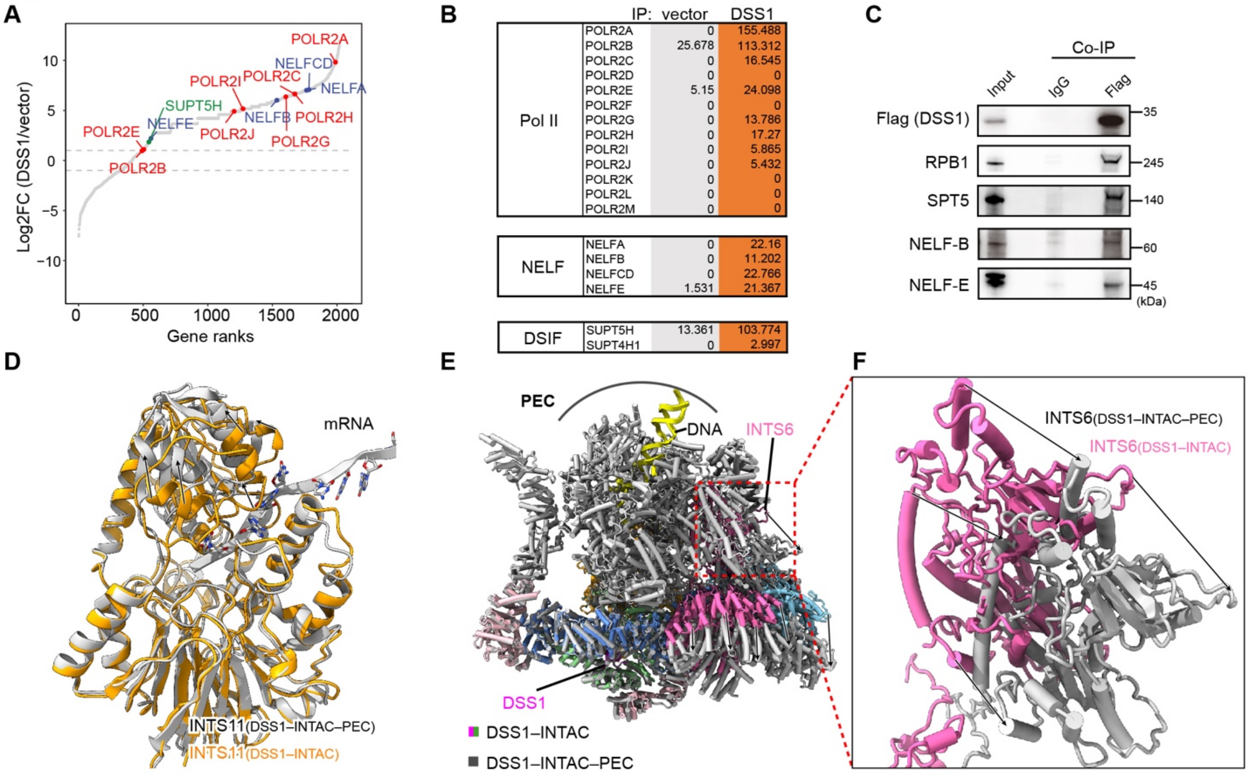

**Fig. S4. DSS1 associates with INTAC in complex with PEC.**

(A) Proteins identified by MS after DSS1 purification were ranked by the enrichment in purification of DSS1 versus vector, with PEC components highlighted. Pol II subunits are labeled in red. NELF subunits are labeled in blue. DSIF subunits are labeled in green. (B) The values of posterior error probability (PEP) score showing the abundance of identified PEC components. (C) Co-IP analysis of Flag (DSS1) followed by western blotting in DSS1–Flag–dTAG cells. (D) Overlaying comparison of INTS11 from the DSS1–INTAC (orange) and DSS1–INTAC–PEC (gray) structures. (E) Overlaying comparison of INTAC from the DSS1–INTAC (colored) and DSS1–INTAC–PEC (gray) structures. The alignment is based on INTS1, INTS2, and INTS7. (F) Close-up view of INTS6 from the DSS1–INTAC (pink) and DSS1–INTAC–PEC (gray) structures.

**Figure S5**

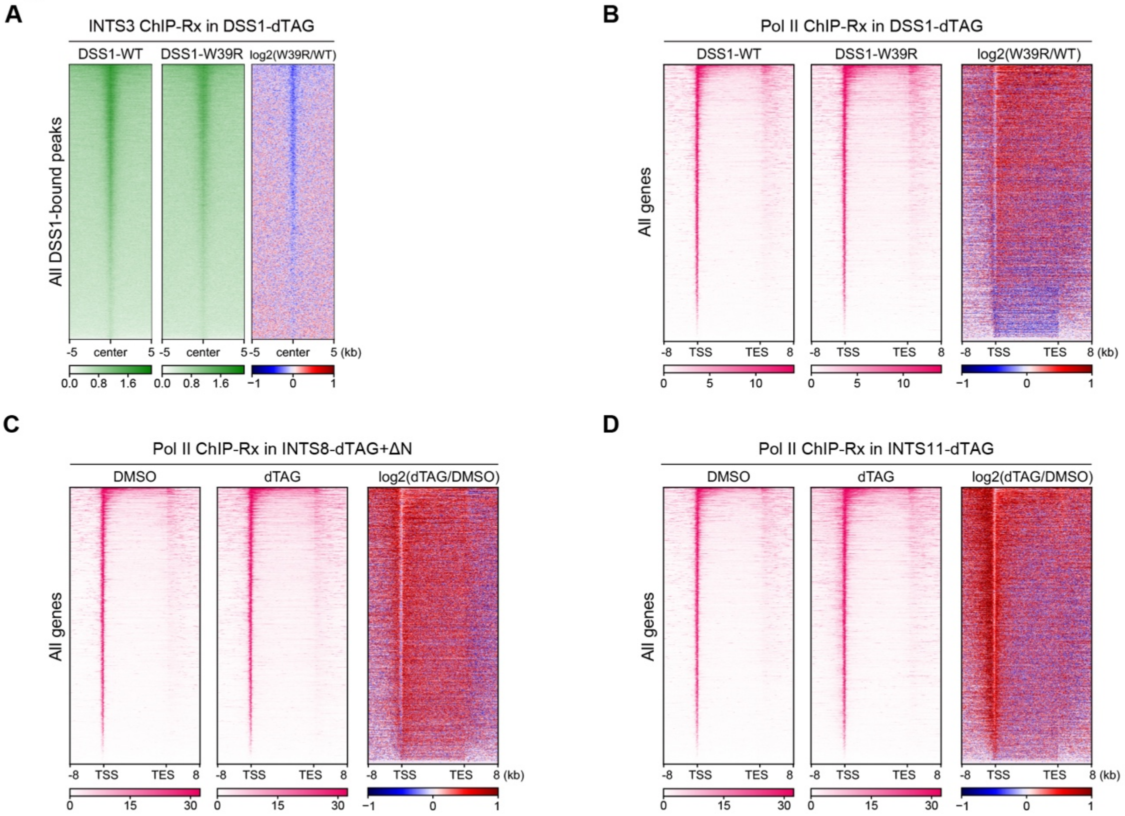

**Fig. S5. DSS1 is required for INTAC function in transcription.**

(A) Heatmaps showing ChIP-Rx signals of INTS3 over 10 kb regions centered on all DSS1-bound peaks after endogenous DSS1 degradation in DSS1–Flag–dTAG cells with the overexpression of DSS1-WT or DSS1-W39R. (B) Heatmaps showing Pol II ChIP-Rx signals of all genes in DSS1–Flag–dTAG cells overexpressing DSS1-WT or DSS1-W39R. (C and D) Heatmaps showing Pol II ChIP-Rx signals of all genes in INTS8-dTAG+ΔN (C) and INTS11-dTAG cells (D) treated with DMSO or dTAG.

**Table S1. Cryo-EM data collection, refinement and validation statistics for DSS1–INTAC and DSS1–INTAC–PEC.**

| | DSS1–INTAC | DSS1–INTAC–PEC |
|---|---|---|
| **Data collection and processing** | | |
| Magnification | 130,000 | 130,000 |
| Voltage (kV) | 300 | 300 |
| Electron exposure (e–/Å$^2$) | 50 | 50 |
| Defocus range (μm) | -1.5 to -2.5 | -1.5 to -2.5 |
| Pixel size (Å) | 1.055 | 1.054 |
| Symmetry imposed | *C1* | *C1* |
| Initial particle images (no.) | 587,397 | 843,523 |
| Final particle images (no.) | 43,790 | 26,822 |
| Map resolution (Å) | 4.1 | 4.6 |
| FSC threshold | 0.143 | 0.143 |
| **Refinement** | | |
| Initial model used (PDB code) | 7YCX | |
| Model resolution (Å) | 4.3 | |
| FSC threshold | 0.5 | |
| Map sharpening *B* factor (Å$^2$) | -124 | |
| Model composition | | |
| Non-hydrogen atoms | 62,554 | |
| Protein residues | 8,285 | |
| Nucleotides | N/A | |
| Ligands | 4 | |
| *B* factors (Å$^2$) | 128.59 | |
| Protein | 128.59 | |
| Nucleotides | N/A | |
| Ligand | 121.11 | |
| R.m.s. deviations | | |
| Bond lengths (Å) | 0.005 | |
| Bond angles (°) | 1.07 | |
| Validation | | |
| MolProbity score | 1.84 | |
| Clashscore | 7.50 | |
| Poor rotamers (%) | 0.05 | |
| Ramachandran plot | | |
| Favored (%) | 93.56 | |
| Allowed (%) | 6.05 | |
| Disallowed (%) | 0.39 | |
| EMDB | | |
| PDB | | |

**Table S2. Detailed information for reagents and materials.**
